# Sites of active gene regulation in the prenatal frontal cortex and their role in neuropsychiatric disorders

**DOI:** 10.1101/2021.09.01.458548

**Authors:** Manuela R. Kouakou, Darren Cameron, Eilis Hannon, Emma L. Dempster, Jonathan Mill, Matthew J. Hill, Nicholas J. Bray

**Affiliations:** MRC Centre for Neuropsychiatric Genetics & Genomics, Division of Psychological Medicine & Clinical Neurosciences, Cardiff University, Cardiff, United Kingdom; University of Exeter Medical School, University of Exeter, United Kingdom

## Abstract

Common genetic variation appears to largely influence risk for neuropsychiatric disorders through effects on gene regulation. It is therefore possible to shed light on the biology of these conditions by testing for enrichment of associated genetic variation within regulatory genomic regions operating in specific tissues or cell types. Here, we have used ATAC-Seq to map open chromatin (an index of active regulatory genomic regions) in bulk tissue, NeuN+ and NeuN− nuclei from the prenatal human frontal cortex, and tested enrichment of SNP heritability for 5 neuropsychiatric disorders (autism spectrum disorder, ADHD, bipolar disorder, major depressive disorder and schizophrenia) within these regions. We observed significant enrichment of SNP heritability for ADHD, major depressive disorder and schizophrenia within open chromatin regions mapped in bulk fetal frontal cortex, and for all 5 tested neuropsychiatric conditions when we restricted these sites to those overlapping histone modifications indicative of enhancers (H3K4me1) or promoters (H3K4me3) in fetal brain. SNP heritability for neuropsychiatric disorders was significantly enriched in open chromatin regions identified in fetal frontal cortex NeuN- as well as NeuN+ nuclei overlapping fetal brain H3K4me1 or H3K4me3 sites. We additionally demonstrate the utility of our mapped open chromatin regions for prioritizing potentially functional SNPs at genome-wide significant risk loci for neuropsychiatric disorders. Our data provide evidence for an early neurodevelopmental component to a range of neuropsychiatric conditions and highlight an important role for regulatory genomic regions active within both NeuN+ and NeuN− cells of the prenatal brain.

## 1. Introduction

The majority of common genetic variants implicated in neuropsychiatric disorders are located in regions of the genome that do not encode proteins (Ripke et al, 2020; Wray et al, 2018; Demontis et al, 2019; Grove et al, 2019; Mullins et al, 2021). These non-coding genomic regions contain *cis-*regulatory elements such as gene promoters, enhancers and silencers that are variably utilised in order to control gene expression in a cell-specific manner (ENCODE Project Consortium, 2012; Roadmap Epigenomics Consortium, 2015). It is possible to map regulatory genomic sites operating in a given tissue or cell-type based on epigenomic features. These include ‘open’, or ‘accessible’, chromatin, which exposes DNA to the transcription factors that modulate gene expression (Tsompana & Buck, 2014). As regulatory regions are believed to contain much of the common genetic component of complex traits, tissues and cell types that are relevant to the etiology of a trait can be delineated by testing for enrichment of trait-associated genetic variation within cell-specific epigenomic features (Finucane et al, 2015).

Several neuropsychiatric disorders are hypothesised to have origins in early brain development (Weinberger et al, 1987; O’Donnell & Meaney, 2017; Courchesne et al, 2020). Consistent with this view, we have recently found that genetic variants associated with altered gene expression (expression quantitative trait loci; eQTL) in the human fetal brain are enriched among risk variants for attention deficit hyperactivity disorder (ADHD), bipolar disorder and schizophrenia (O’Brien et al, 2018). Similarly, de la Torre-Ubieta and colleagues (2018) mapped genomic regions of open chromatin in the germinal zone and cortical plate of the fetal cerebral cortex, reporting significant enrichment of genetic variation associated with ADHD, depressive symptoms, neuroticism and schizophrenia within sites defined as preferentially accessible in the germinal zone. In the present study, we used the Assay for Transposase-Accessible Chromatin with high-throughput sequencing (ATAC-Seq; Buenrostro et al, 2013) to map regions of open chromatin in the human second trimester frontal cortex, finding these to be enriched for genetic variation associated with a range of neuropsychiatric disorders, particularly when histone modifications indicative of enhancers or promoters were also considered. In addition, we provide the first maps of open chromatin in NeuN+ (neuron-enriched) and NeuN− (neuron-depleted) nuclei from the human fetal brain, and demonstrate use of our open chromatin maps for prioritizing potentially functional SNPs at genome-wide significant risk loci for neuropsychiatric disorders.

## 2. Methods

### 2.1 Samples

Open chromatin annotations were derived from 3 fresh human fetal frontal cortex samples from the second trimester of gestation (16, 18 and 19 post-conception weeks). This number of biological replicates is consistent with other epigenomic studies of its kind (e.g. de la Torre-Ubieta et al, 2018) and meets current ENCODE ATAC-Seq standards for genomic annotation, which stipulate at least 2 independent replicates. Fetal cortex samples were acquired from the MRC-Wellcome Trust Human Developmental Biology Resource (HDBR) (http://www.hdbr.org/) in Hibernate-E media (ThermoFisher Scientific). All samples were obtained through elective terminations of pregnancy, with consent from female donors, and were of normal karyotype (2 female and 1 male). Ethical approval for the collection and distribution of fetal material for scientific research was granted to the HDBR by the Royal Free Hospital research ethics committee (reference 08/H0712/34) and NRES Committee North East - Newcastle & North Tyneside (reference 08/H0906/21+5). The left frontal cortex from each foetus was dissected and dounce homogenized on ice to produce a single cell suspension. Prior to nuclei isolation, aliquots of 1 to 10 million cells were stored at −80°C in cryovials containing 1ml Hibernate-E media, supplemented with 6% dimethyl sulfoxide (DMSO), in Nalgene^®^ Mr Frosty containers.

### 2.2 Nuclei isolation and Assay for Transposase-Accessible Chromatin in bulk tissue

Two technical replicates were processed from each of the 3 bulk frontal cortex samples. Nuclei were isolated from cryopreserved cell suspensions by first centrifuging cells at 600g for 5 mins and resuspending them in ice-cold cell lysis buffer (sucrose 0.25M; KCl 25mM, MgCl2 5mM, Tris-Cl 10mM, dithiothereitol 1mM, 0.1% Triton) for 15 mins. Nuclei were then pelleted by centrifuging at 1000g for 8 mins and resuspended in storage buffer (sucrose 0.25M, MgCl2 5mM, Tris-Cl 10mM, BSA 0.1%) by gentle pipetting for 10 secs. The nuclei suspension was then left on ice for 10 mins and passed through a 21G needle 10 times to thoroughly resuspend nuclei. Nuclei were then visualized under an optical microscope for quality control and counting. For each technical replicate, 50,000 nuclei were incubated in the transposition reaction mix (20μl nuclease-free water; 25μl 2X Tagment DNA Buffer, Illumina Cat#FC-121-1030; 5μl Tn5 Transposase, Illumina Cat#FC-121-1030) at 37°C for 30 minutes, as described by Buenrostro et al (2013).

### 2.3 Fluorescence-activated nuclei sorting (FANS)

Nuclei were isolated from cryopreserved frontal cortex cell suspensions as described above. To block non-specific antibody binding, the resulting nuclei pellet was resuspended in buffer containing sucrose 0.25 M, MgCl2 5mM, Tris-Cl 10mM and BSA 1%, and incubated for 30 minutes on ice. Nuclei were visualized under an optical microscope for quality control and counting. The samples were then centrifuged at 400g for 8 mins and the nuclei pellet re-suspended in FANS buffer (0.5% BSA in DPBS).

Before immunostaining, 100μl of the sample was transferred to a new tube and used as unstained control in the FANS analysis. The remaining sample was incubated with mouse anti-neuronal nuclei (NeuN) Alexa Fluor 488 conjugated monoclonal antibody (MerkMillipore Cat# MAB377X), at a 1:2000 dilution, and rotated at 4°C in the dark for 60 mins. After immunostaining, samples were washed 3 times in FANS buffer (400g for 5 min) to remove excess antibody. DAPI was then added to a final concentration of 1μg/ml and the nuclei suspension was filtered through a 35μm cell strainer to remove nuclei clumps and prevent clogging of the cytometer. DAPI positive neuronal (NeuN+) and non-neuronal (NeuN−) nuclei were sorted into tubes pre-coated with 5% BSA using a FACSAria flow cytometer (BD Biosciences) equipped with a 100μm nozzle. For both flow analysis and FANS, exclusion of debris using forward and side scatter pulse area parameters (FSC-A and SSC-A) were gated first, followed by exclusion of aggregates using pulse width (FSC-W and SSC-A) and DAPI-positive nuclei before gating populations based on NeuN fluorescence (Supplementary Figure S1). Sorted NeuN+ and NeuN− nuclei were incubated in transposition reaction mix and PCR amplified as for bulk tissue. Sufficient nuclei were recovered to perform two technical replicates for 2 of the 3 frontal cortex samples and 1 technical replicate for the other, resulting in 5 separate transposase reactions for NeuN+ nuclei (mean number of nuclei per reaction = 41,216) and 5 separate transposase reactions for NeuN− nuclei (mean number of nuclei per reaction = 44,066).

### 2.4 PCR amplification, size selection and sequencing of ATAC-Seq libraries

Transposase reactions were initially amplified by 8 cycles of PCR using primers described by Buenrostro et al (2013) and 2X NEBNext Q5 HotStart HiFi Master Mix (New England Biolabs, Cat# M0543). Amplicons of 175 -250bp were size-selected using agarose gel electrophoresis on BluePippin 2% gel cassettes (Sage Science) and further amplified through 7 additional PCR cycles. Fragment size analysis and quantification of ATAC-seq libraries were performed using an Agilent Technologies 2100 Bioanalyzer and high sensitivity DNA kit and the Qubit dsDNA HS Assay Kit (ThermoFisher Cat#32854). ATAC-Seq libraries were sequenced on an Illumina HiSeq 4000 to a depth of at least 100 million paired-end 75bp reads per library.

### 2.5 ATAC-Seq data analysis

Sequencing quality was confirmed using FastQC (http://www.bioinformatics.babraham.ac.uk/projects/fastqc). Reads were aligned to the GRCh37 (hg19) human genome reference sequence using Bowtie2 v2.2.9 (Langmead et al, 2009) following adapter trimming. This produced a SAM file for each replicate which was then converted into a coordinate-sorted BAM file of paired-end reads with Samtools v1.5 (Li et al, 2009). Reads that mapped to more than one locus or were PCR duplicates were excluded. This yielded >90 million uniquely mapped, non-duplicated paired-end reads per technical replicate.

We first down-sampled reads from the 6 BAM files (2 technical replicates for each of the 3 biological replicates) for bulk fetal frontal cortex to the lowest read count of any technical replicate and merged these using Samtools v1.5 (Li et al, 2009) to produce a large, single down-sampled BAM file representing all samples. Open chromatin peaks (FDR < 0.01) were identified in the down-sampled bulk tissue BAM file using MACS2 (Zhang et al, 2008), while artefact regions (defined as “Blacklist” regions by ENCODE; https://www.encodeproject.org/files/ENCFF001TDO/) with excessive unstructured anomalous read mapping (e.g. regions of centromeres, telomeres and satellite repeats) were excluded. We then merged the 2 technical replicate BAM files for each biological replicate to obtain 3 biological replicate BAM files. MACS2 (Zhang et al, 2008) was used to identify peaks (FDR < 0.01) in each biological replicate and DiffBind v2.16.0 (Stark & Brown, 2011) to determine peaks observed in at least 2 of the 3 biological replicates. To generate a set of high confidence bulk fetal frontal cortex OCRs for our main analyses, we selected peaks (FDR < 0.01) in the single down-sampled BAM file that intersected with the (FDR < 0.01) peaks observed in at least 2 of the 3 biological replicates. The 5 technical replicate BAM files for NeuN+ and NeuN− nuclei were similarly down-sampled and merged before identifying open chromatin peaks (FDR < 0.01) using MACS2 (Zhang et al, 2008). To generate subsets of bulk fetal frontal cortex OCRs that could be attributed to the NeuN+ and / or NeuN− fractions, we selected those high confidence bulk peaks that intersected with the FDR < 0.01 peaks identified in the merged BAM files for both NeuN+ and NeuN− nuclei.

### 2.6 Overlap between open chromatin regions identified in different tissues

ChIPpeakAnno (Zhu et al, 2010) was used to determine the numbers of bulk fetal frontal OCRs overlapping adult frontal cortex OCRs (Hoffman et al, 2019) and OCRs identified in the germinal zone / cortical plate of the fetal cortex (de la Torre-Ubieta et al, 2018). Consensus OCRs identified by ATAC-Seq in bulk adult frontal cortex (CMC_ATACSeq_consensusPeaks.bed) were downloaded from https://www.synapse.org/#!Synapse:syn18134202 and converted to GRCh37 coordinates using https://genome.ucsc.edu/cgi-bin/hgLiftOver. OCRs identified by ATAC-Seq in the germinal zone / cortical plate of the fetal cortex were downloaded from https://www.ncbi.nlm.nih.gov/geo/query/acc.cgi?acc=GSE95023.

### 2.7 Bioinformatic annotation of open chromatin regions

We tested for overlap between bulk fetal frontal cortex OCRs and chromatin states that have been defined in independent samples of (bulk) human fetal brain and regions of the adult human brain on the basis of other epigenomic assays (histone modification ChIP-Seq, transcription factor ChIP-Seq and DNAse-Seq; Libbrecht et al, 2019). Annotation maps were downloaded from https://noble.gs.washington.edu/proj/encyclopedia/interpreted/ and the R package ChIPseeker (Yu et al, 2015) used to simulate 5,000 sets of regions that matched the genomic distribution of the bulk fetal frontal cortex OCRs in terms of chromosome distribution and region sizes. Taking all regions annotated to the same chromatin state for each sample in turn, we counted the number of overlapping peaks with the simulated set of OCR peaks. An overlap was defined as >50% of the OCR peak intersecting an annotated region. The mean number of intersecting regions calculated across all simulations was compared to the true overlap to calculate a fold-enrichment statistic. One-sided empirical *P*-values for both over- and under enrichment were calculated separately as the number of simulations with a more extreme overlap divided by the total number of simulations, adding 1 to both the denominator and numerator. Two-sided *P*-values were calculated as a) the sum of the one-sided *P*-value for over-enrichment and 1 minus the one-sided *P*-value for under-enrichment, if the fold-change was >1, or b) the sum of one-sided *P*-value for under-enrichment and 1 minus the one-sided *P-*value for over-enrichment, if the fold-change was <1.

We tested for enrichment of transcription factor binding motifs within high confidence fetal frontal cortex OCRs using HOMER (http://homer.ucsd.edu/homer/ngs/peakMotifs.html). Enrichment of biological process Gene Ontology (GO) annotations for genes with fetal frontal cortex OCRs within 30kb upstream and 100bp downstream of their TSS was tested using g:Profiler (Raudvere et al, 2019), correcting for multiple testing using the default g:SCS algorithm. We tested for enrichment of previously identified genome-wide significant (*P* < 5 × 10^−8^) eQTL (O’Brien et al, 2018) and mQTL (Hannon et al, 2016) operating in the human fetal brain within fetal frontal cortex OCRs using GARFIELD (Iotchkova et al, 2019), a method that controls for minor allele frequency, GC-content, linkage disequilibrium and local gene density.

### 2.8 Histone modification datasets from human fetal brain

Genomic regions marked by H3K4me1 or H3K4me3 in a bulk human fetal brain sample of similar gestational age to those used in this study (sample E082; female, 17 post-conception weeks) were downloaded as BED files from the Roadmap Epigenomics Project (https://egg2.wustl.edu/roadmap/data/byFileType/peaks/consolidated/narrowPeak/). BEDTools v2.26.0 intersect and merge (Quinlan & Hall, 2010) were used to identify and merge H3K4me1 and H3K4me3 sites overlapping the OCRs that we identified in nuclei from the fetal frontal cortex.

### 2.9 Testing enrichment of SNP heritability for neuropsychiatric disorders in open chromatin regions

Stratified LD score regression (SLDSR; Finucane et al, 2015) was used to test for enrichment of SNP heritability for neuropsychiatric disorders within fetal brain OCRs, H3K4me1 sites, H3K4me3 sites and OCRs overlapping these sites. Peaks were expanded by 500bp on each side prior to SLDSR, as recommended by Finucane et al (2015). Summary statistics from large-scale GWAS of 5 neuropsychiatric disorders (ADHD [Demontis et al, 2019], autism spectrum disorder [Grove et al, 2019], bipolar disorder [Mullins et al, 2021], major depressive disorder [Wray et al, 2018] and schizophrenia [Ripke et al, 2020]) and two control traits with similar sample sizes (height [Lango Allen et al, 2010] and blood triglyceride levels [Teslovich et al, 2010]) were tested (Supplementary Table S6). Fold-enrichment estimates are the proportion of SNP heritability for each trait explained by SNPs within the annotation divided by the proportion of genome-wide SNPs within the annotation. Each annotation was also assessed against the baseline model provided by Finucane et al (consisting of 53 genomic annotations including coding regions, promoters, enhancers and conserved regions) and resulting Z-scores used to calculate two-tailed *P*-values. We more conservatively state the Z-score *P*-values, rather than raw fold-enrichment *P*-values, in the text and figures, highlighting those that survive Bonferroni correction for the 7 tested traits (i.e. *P* < 0.0071).

### 2.10 Identification of SNPs within open chromatin regions in strong linkage disequilibrium with index SNPs from GWAS of neuropsychiatric disorders

We used GARFIELD (Iotchova et al, 2019) to identify SNPs within high confidence bulk fetal frontal cortex OCRs that exhibit genome-wide significant (*P* < 5 × 10^−8^) association with any of the 5 tested neuropsychiatric disorders in the associated GWAS. We used LDpair within the NIH national Cancer Institute LDlink (https://ldlink.nci.nih.gov/) to further identify those OCR SNPs that were in strong linkage disequilibrium (r2 > 0.8) with the most significant (index) SNP at each genome-wide significant risk locus in the CEU population. To further prioritise potentially functional OCR SNPs, we identified those for which there was evidence that they act as eQTL in the human fetal brain (*P* < 5 × 10^−5;^ O’Brien et al, 2018), and used LDpair to determine the r2 between them and the most significant eQTL for the implicated transcript in the CEU population.

## 3. Results

We used ATAC-Seq (Buenrostro et al, 2013) to map open chromatin regions (OCRs) in the frontal cortex of 3 human fetuses from the second trimester of gestation. We first profiled OCRs within unsorted nuclei from these samples (‘bulk’ frontal cortex analyses), identifying 88,501 high confidence OCRs using stringent peak calling criteria (see Methods), and observing strong correlations in the number of reads within peaks between biological replicates (median r = 0.83; *P* < 2.2 × 10^−16^; Supplementary Figure S2). The genomic distribution of peaks approximated that of OCRs identified in other tissues (Song et al, 2011; Bryois et al, 2018), with 28% within 5kb of a transcription start site (TSS) and 38% within gene bodies. Exemplar fetal frontal cortex OCRs, located at the TSS and first intron of the gene encoding the neurogenic transcription factor NeuroD1, are shown in Figure 1a. Identified OCRs were prominently enriched in genomic regions annotated as active promoters and enhancers (Libbrecht et al, 2019) on the basis of histone modifications and other epigenomic features in independent bulk human fetal brain samples (54.3- and 6.1-fold enrichment respectively; *P* < 0.0002), while showing a general depletion in regions annotated as quiescent or repressed in the fetal and adult brain (Figure 1b; Supplementary Table S1). Fetal frontal cortex OCRs were highly enriched for the CTCF binding motif (11.8-fold enrichment, *P* = 1 × 10^−5440^), with notable enrichment of motifs for neural transcription factors such as Olig2 (2.2-fold enrichment, *P* = 1 × 10^−1067^) and Oct6 (2.1-fold enrichment, *P* = 1 × 10^−449^) also observed. Consistent with roles in gene regulation, fetal frontal cortex OCRs were enriched for both expression quantitative trait loci (eQTL; 2.1-fold enrichment, *P* = 3.47 × 10^−19^) and methylation quantitative trait loci (mQTL, 2.3-fold enrichment, *P* = 1.88 × 10^−23^) identified in the human fetal brain (O’Brien et al, 2018; Hannon et al, 2016).

**Figure 1.**
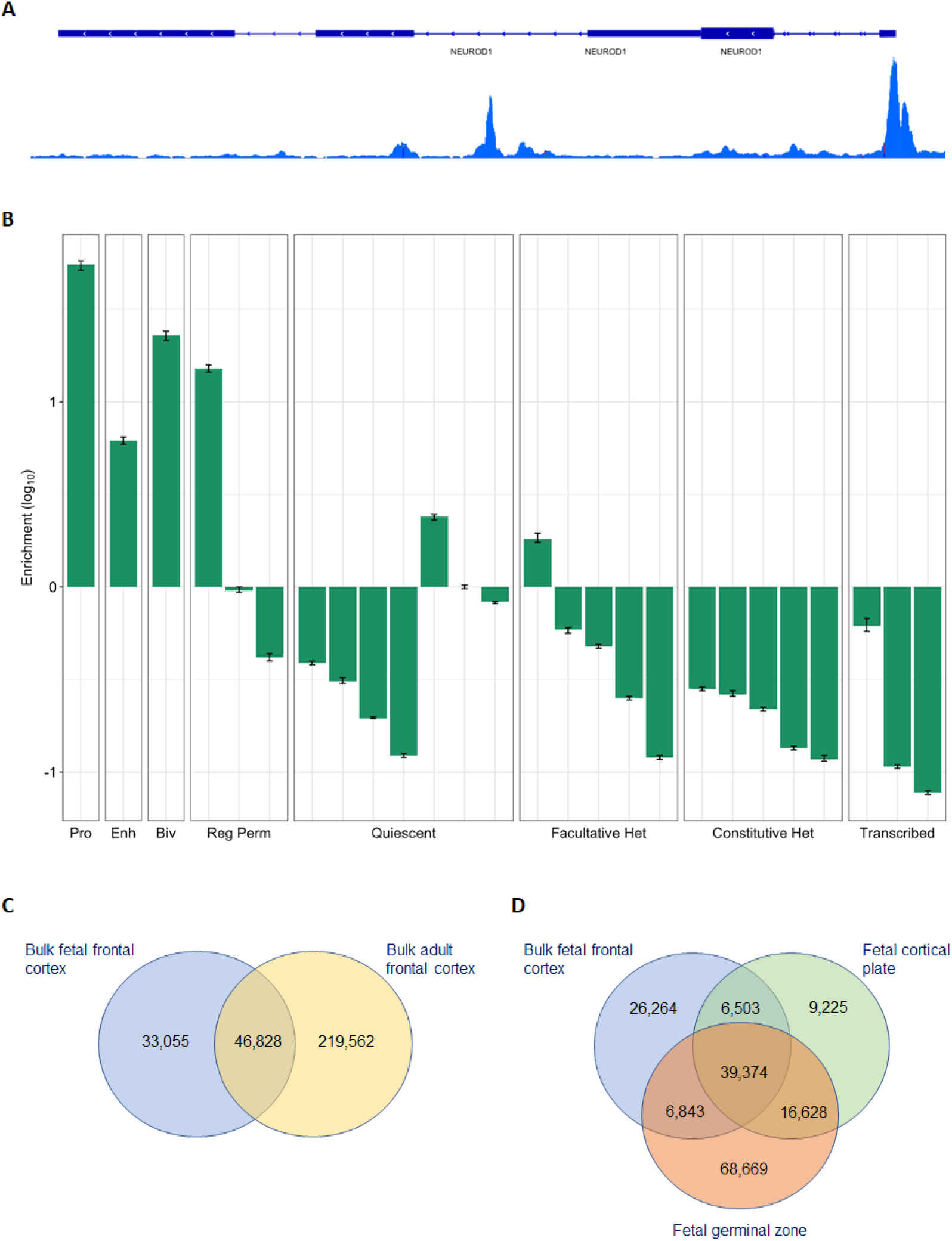
Characteristics of high confidence bulk fetal frontal cortex open chromatin regions. A) Combined reads from the 3 bulk fetal frontal cortex samples showing a high confidence open chromatin region at the transcription start site of the *NEUROD1* gene. B) Log10 enrichments of fetal frontal cortex open chromatin regions within functional genomic annotations defined by Libbrecht et al (2019) in bulk human fetal brain based on histone modifications and other epigenomic features. Pro = Promoter; Enh = Enhancer; Biv = Bivalent: regulatory element with marks of both activation (such as H3K27ac) and repression (H3K27me3); Reg Perm = RegPermissive: region with weak marks of regulatory activity such as H3K4me1 or DNase hypersensitivity; Quiescent: Inactive region; Facultative Het: heterochromatin marking regions of cell type-specific repression, characterized by the histone modification H3K27me3 (also known as Polycomb-repressed heterochromatin); Constitutive Het: heterochromatin marking permanently silent regions, characterized by the histone modification H3K9me3; Transcribed: transcribed genic region. C) Overlap between open chromatin regions identified in bulk fetal frontal cortex and bulk adult frontal cortex (Hoffman et al, 2019). d) Overlap between open chromatin regions identified in bulk fetal frontal cortex and those identified in cortical plate and germinal zone of the fetal cortex in the study of de la Torre-Ubieta et al (2018). Note that the minimum number of overlapping peaks is calculated, and therefore the summed number of open chromatin regions will usually be smaller than the total number in each set.

Approximately 60% of the OCRs we identified in fetal frontal cortex overlapped OCRs observed in adult human frontal cortex (Hoffman et al, 2019; Figure 1c); genes with non-overlapping ‘fetal-specific’ OCRs within 30kb upstream of their TSS were most significantly enriched for the Gene Ontology (GO) term ‘nervous system development’ (*P corrected* = 6.7 × 10^−9^), while the larger number of genes with ‘adult-specific’ OCRs within 30kb upstream of their TSS were highly enriched for the GO terms ‘signaling’ (*P corrected* = 2.6 × 10^−60^) and ‘cell communication’ (*P corrected* = 1.2 × 10^−59^). There was also substantial overlap with OCRs previously observed in both the germinal zone and cortical plate of the human fetal brain (de la Torre-Ubieta et al, 2018; Figure 1d), although we note that ~33% of our high-confidence fetal cortex OCRs were not identified in either of these regions.

We tested for enrichment of single nucleotide polymorphism (SNP) heritability for 5 major neuropsychiatric disorders (ADHD, autism spectrum disorder [ASD], bipolar disorder, major depressive disorder and schizophrenia) within bulk fetal frontal cortex OCRs using stratified linkage disequilibrium score regression (SLDSR; Finucane et al, 2015), controlling for general genomic annotations (e.g. coding regions, promoters, enhancers and conserved regions) included in the baseline model to obtain Z-score *P*-values. We observed significant enrichment of SNP heritability for ADHD (5.6-fold enrichment, Z-score *P* = 0.009), major depressive disorder (4.4-fold enrichment, Z-score *P* = 0.024) and schizophrenia (7.2-fold enrichment, Z-score *P* = 1.2 × 10^−11^) within these regions, the latter surviving Bonferroni correction for the 7 tested traits (Supplementary Table S2). In contrast, we observed no such enrichment of SNP heritability for either of the control traits (triglyceride levels or height) when baseline annotations were accounted for (Z-score *P* > 0.1 for both). The level of enrichment of schizophrenia risk variation in fetal frontal cortex OCRs approximates that reported in adult frontal cortex OCRs (Bryois et al, 2018), with 3.3% of SNPs in fetal brain OCRs accounting for 23.9% of SNP heritability for the condition.

A recent study (Schork et al, 2019) reported enrichment of SNP heritability for a broad neuropsychiatric phenotype encompassing ADHD, affective disorder, anorexia, ASD, bipolar disorder and schizophrenia within H3K4me1 and H3K4me3 sites (histone modifications indicative of poised or active enhancers and promoters, respectively) identified by the RoadMap Epigenomics Mapping Consortium (2015) in bulk human fetal brain tissue. Consistent with these findings, when we restricted our bulk fetal frontal cortex OCRs to those overlapping either fetal brain H3K4me1 or H3K4me3 sites identified by the RoadMap Epigenomics Mapping Consortium, we observed strong enrichment of SNP heritability for all 5 tested neuropsychiatric disorders (but not for the two control traits), with Z-score *P*-values surviving Bonferroni correction in each case (Figure 2; Supplementary Table S2). With the exception of enrichment of SNP heritability for major depressive disorder in fetal brain OCRs overlapping H3K4me3 sites, enrichments within fetal brain OCRs overlapping H3K4me1 or H3K4me3 regions were consistently higher than within fetal brain OCRs or histone modification sites alone (Figure 3), highlighting the value of additional epigenomic annotations to define regulatory regions of the genome.

**Figure 2.**
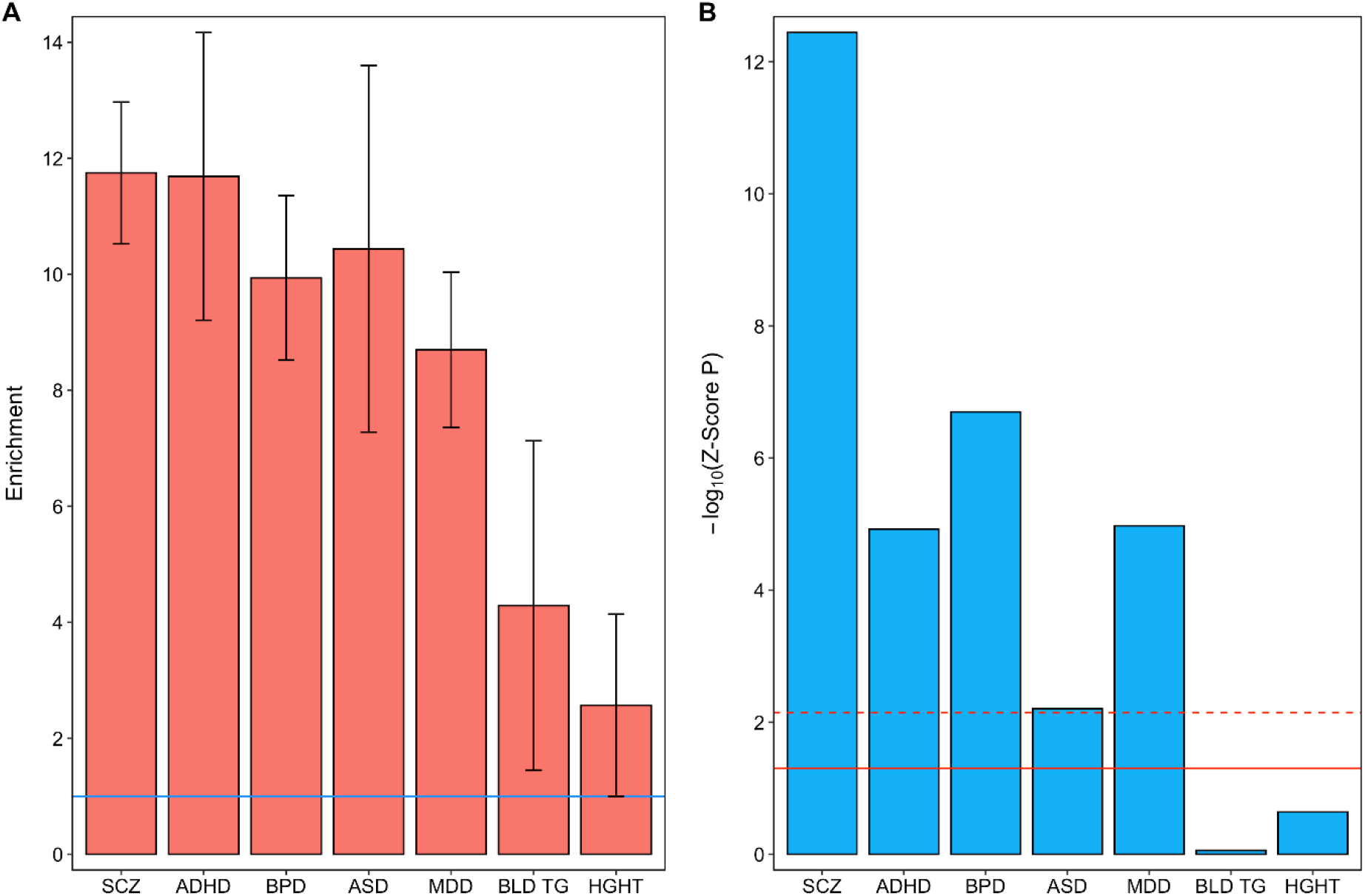
Partitioned heritability for 5 neuropsychiatric disorders and 2 control traits within open chromatin regions identified in bulk fetal frontal cortex overlapping fetal brain H3K4me1 sites. A) Fold-enrichment estimates of SNP heritability for each trait (the proportion of SNP heritability explained by SNPs within the annotation divided by the proportion of genome-wide SNPs within the annotation). Error bars represent standard error. The solid horizontal line indicates no enrichment. B) −Log_10_ Z-score *P*-values (two-tailed) for enrichment of SNP heritability, controlling for general genomic annotations included in the baseline model of Finucane and colleagues (2015). The solid horizontal line indicates the Z-score *P*-value 0.05 threshold; the dashed horizontal line indicates the threshold for Z-score *P*-values surviving Bonferroni correction for 7 tested traits. SCZ = schizophrenia; ADHD = attention deficit hyperactivity disorder; BPD = bipolar disorder; MDD = major depressive disorder; ASD = autism spectrum disorder; BLD TG = blood triglyceride levels; HGHT = height.

**Figure 3.**
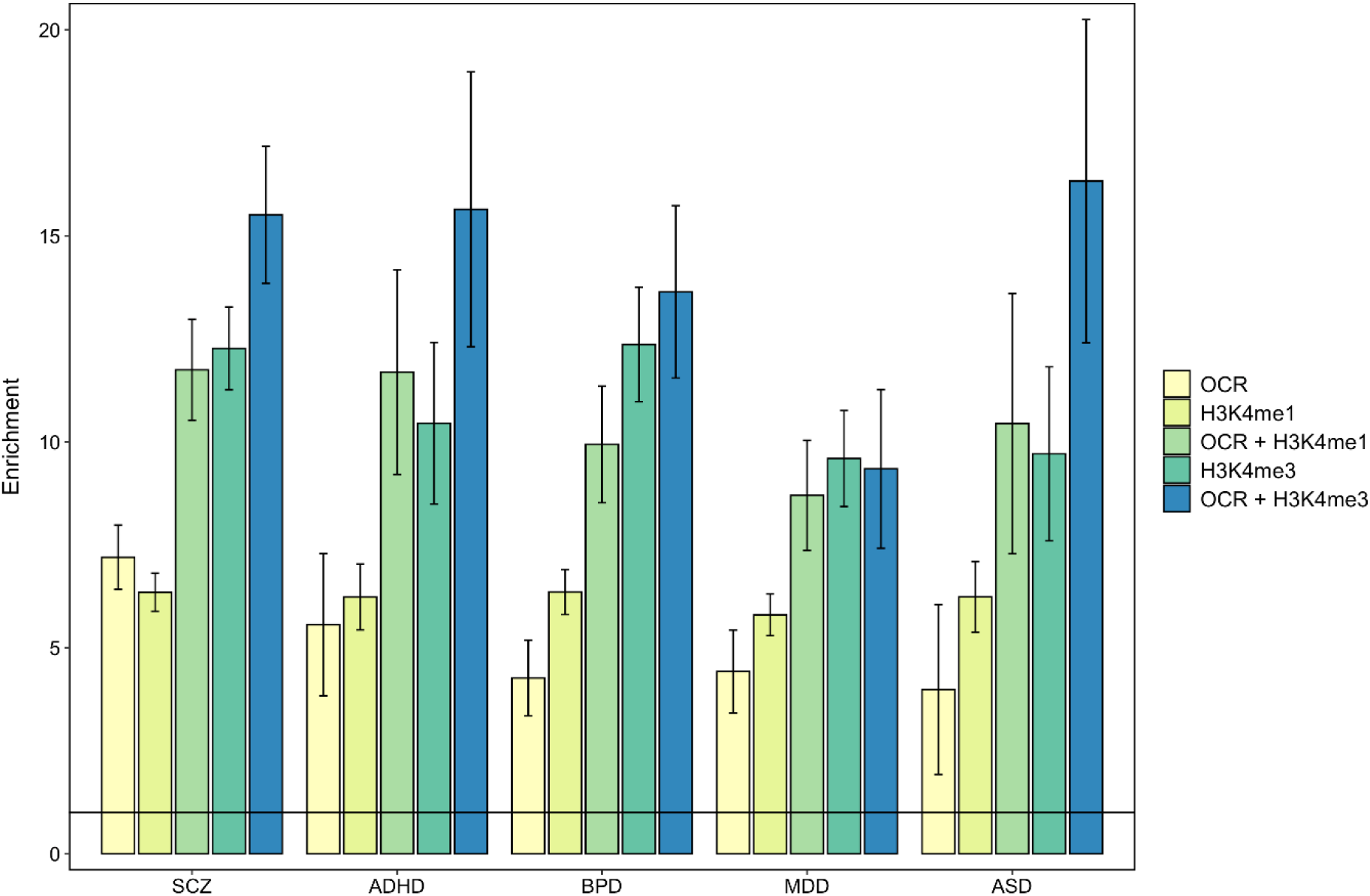
Enrichment of SNP heritability for 5 neuropsychiatric disorders within open chromatin regions identified in bulk fetal frontal cortex (OCR), within previously identified sites of histone modification in the human fetal brain (H3K4me1 and H3K4me3) and within bulk fetal frontal cortex open chromatin regions overlapping each histone modification (OCR + H3K4me1 and OCR + H3K4me3). Fold-enrichment estimates of SNP heritability for each trait is the proportion of SNP heritability explained by SNPs within the annotation divided by the proportion of genome-wide SNPs within the annotation. The solid horizontal line indicates no enrichment. Error bars represent standard error. SCZ = schizophrenia; ADHD = attention deficit hyperactivity disorder; BPD = bipolar disorder; MDD = major depressive disorder; ASD = autism spectrum disorder.

In the adult brain, common genetic risk for schizophrenia has been reported to be primarily mediated through OCRs in NeuN+ (neuronal), rather than NeuN− (non-neuronal), nuclei (Fullard et al, 2018). To explore the cellular basis of genetic risk for neuropsychiatric disorders in the prenatal brain, we determined which of our fetal frontal cortex OCRs could be confidently attributed to NeuN+ and NeuN− fractions by fluorescence-activated sorting nuclei from the same 3 fetal frontal cortex samples. This identified 30,162 high confidence bulk fetal frontal cortex OCRs that were also observed in sorted NeuN+ (neuron-enriched) nuclei (at FDR < 0.01) and 37,576 such OCRs that were also observed in sorted NeuN− (neuron-depleted) nuclei (at FDR < 0.01), of which 27,528 OCRs overlapped both NeuN+ and NeuN− fractions (Figure 4a). OCRs (FDR < 0.01) specific to NeuN+ nuclei included sites at the proximal promoter of *GABRB3* (−141bp from the TSS) and in intron 1 of the neuronal marker *MAP2* (+1614bp from the TSS). OCRs (FDR < 0.01) identified in NeuN− but not NeuN+ nuclei include sites at the proximal promoter of the gene encoding the oligodendrocyte precursor cell marker OLIG2 (−139bp from the TSS), and sites at the proximal promoters of the genes encoding the neural progenitor / radial glia cell markers nestin (*NES*; −399bp from the TSS), SOX9 (−46bp from the TSS) and SOX2 (−63bp from the TSS; Figure 4b), consistent with successful depletion of non-neuronal nuclei in the NeuN+ fraction.

**Figure 4.**
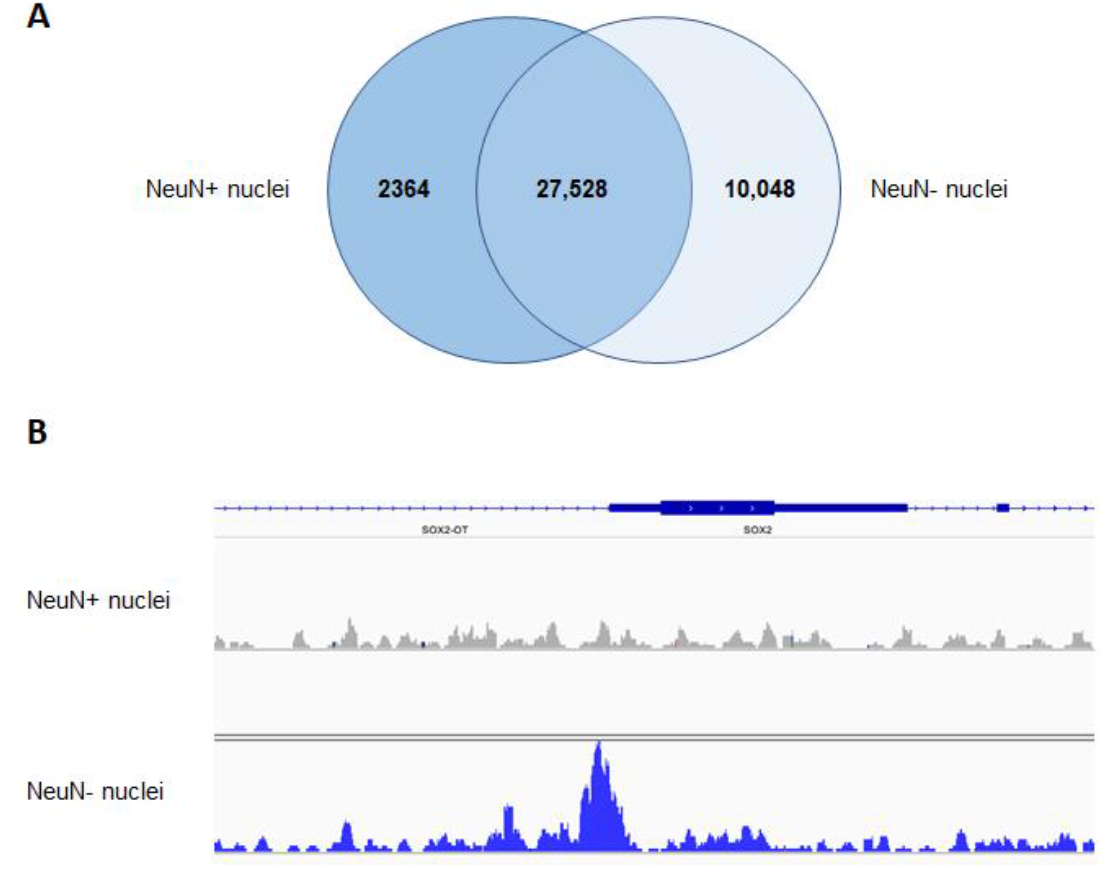
High confidence open chromatin regions identified in NeuN+ and NeuN− nuclei from the fetal frontal cortex. A) Overlap between open chromatin regions identified in fetal frontal cortex NeuN+ and NeuN− nuclei. B) Combined reads from the 3 fetal NeuN+ samples and 3 fetal NeuN− samples showing a high confidence open chromatin peak at the transcription start site of the SOX2 gene in NeuN− but not NeuN+ nuclei.

We observed significant enrichment of SNP heritability for schizophrenia in OCRs identified in fetal frontal cortex NeuN+ nuclei, which survived correction for multiple testing (6.5-fold enrichment, Z-score *P* = 4.16 × 10^−4^). Moreover, SNP heritability for schizophrenia, bipolar disorder and major depressive disorder was enriched at Bonferroni-corrected significance in NeuN+ OCRs overlapping either fetal H3K4me1 sites (Figure 5) or H3K4me3 sites, with nominally significant enrichments observed for ASD in NeuN+ OCRs overlapping H3K4me1 sites and for ADHD in NeuN+ OCRs overlapping H3K4me3 sites (Supplementary Table S3). Within fetal frontal cortex NeuN− nuclei, we observed enrichment of SNP heritability for schizophrenia (8.14-fold enrichment, Z-score *P* = 1.2 × 10^−7^) and ADHD (8.69-fold enrichment, Z-score *P* = 1.54 × 10^−4^) at Z-score *P*-values surviving Bonferroni correction, and for bipolar disorder at nominal significance (5.82-fold enrichment, Z-score *P* = 0.01). For fetal frontal cortex NeuN− nuclei overlapping either fetal H3K4me1 (Figure 6) or H3K4me3 sites, we observed significant enrichment of SNP heritability for all 5 tested neuropsychiatric disorders, each surviving correction for multiple testing (Supplementary Table S4).

**Figure 5.**
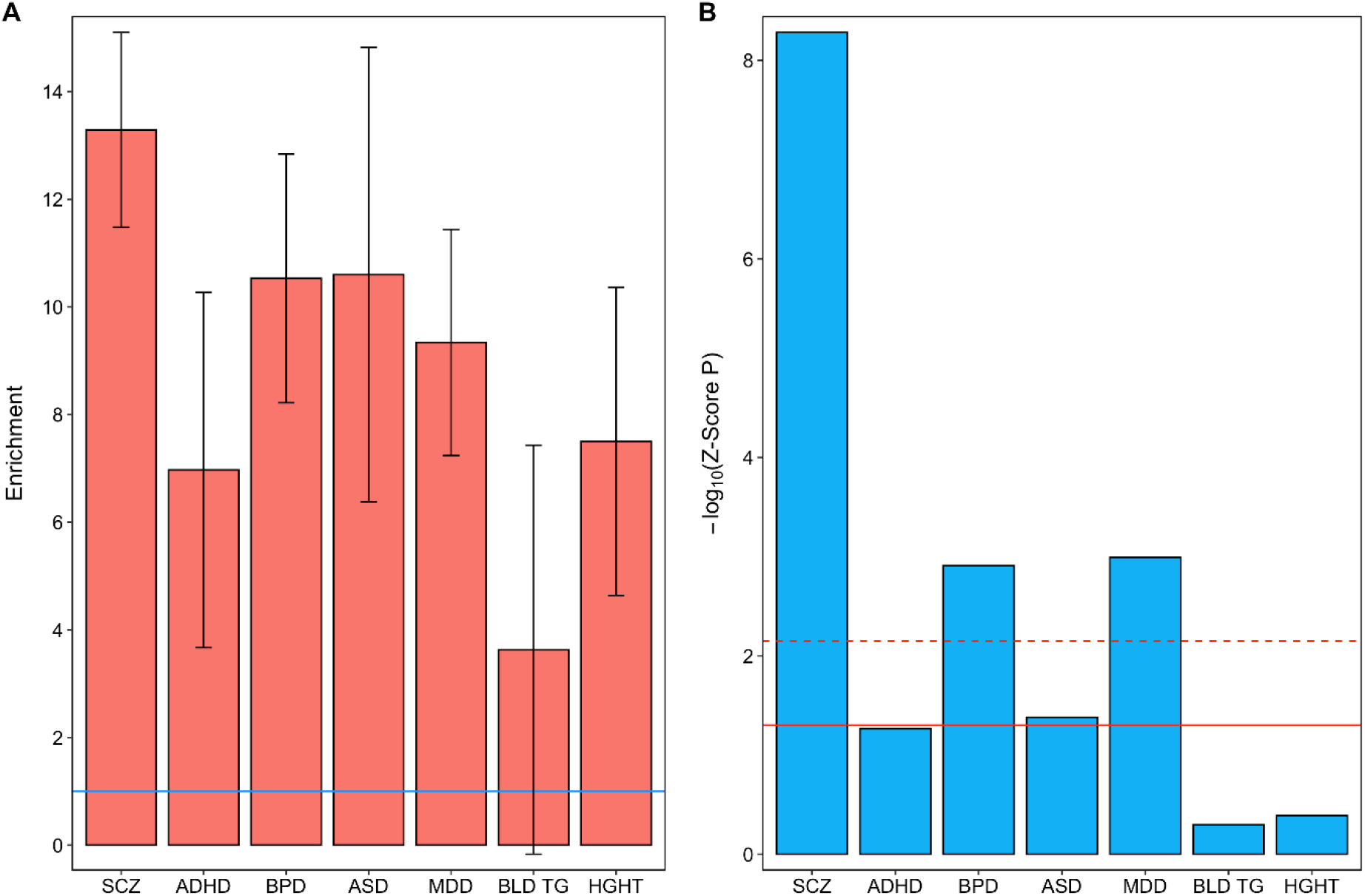
Partitioned heritability for 5 neuropsychiatric disorders and 2 control traits within high confidence open chromatin regions observed in fetal brain NeuN+ nuclei. A) Fold-enrichment estimates of SNP heritability for each trait (the proportion of SNP heritability explained by SNPs within the annotation divided by the proportion of genome-wide SNPs within the annotation). Error bars represent standard error. The solid horizontal line indicates no enrichment. B) -Log_10_ Z-score P-values (two-tailed) for enrichment of SNP heritability, controlling for general genomic annotations included in the baseline model of Finucane and colleagues (2015). The solid horizontal line indicates the Z-score P-value 0.05 threshold; the dashed horizontal line indicates the threshold for Z-score P-values surviving Bonferroni correction for 7 tested traits. SCZ = schizophrenia; ADHD = attention deficit hyperactivity disorder; BPD = bipolar disorder; MDD = major depressive disorder; ASD = autism spectrum disorder; BLD TG = blood triglyceride levels; HGHT = height.

**Figure 6.**
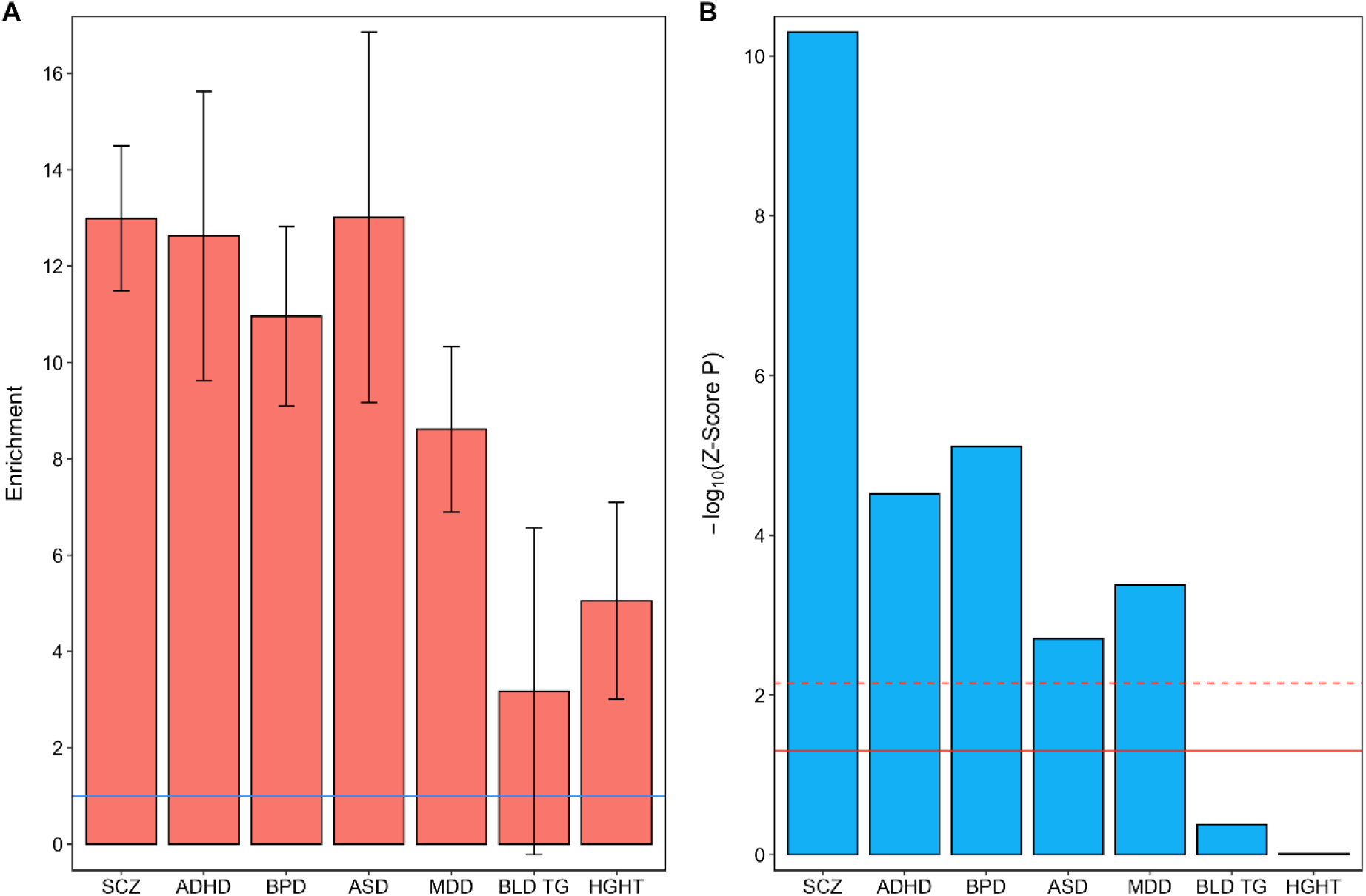
Partitioned heritability for 5 neuropsychiatric disorders and 2 control traits within high confidence open chromatin regions observed in fetal brain NeuN− nuclei. A) Fold-enrichment estimates of SNP heritability for each trait (the proportion of SNP heritability explained by SNPs within the annotation divided by the proportion of genome-wide SNPs within the annotation). Error bars represent standard error. The solid horizontal line indicates no enrichment. B) -Log_10_ Z-score *P*-values (two-tailed) for enrichment of SNP heritability, controlling for general genomic annotations included in the baseline model of Finucane and colleagues (2015). The solid horizontal line indicates the Z-score *P*-value 0.05 threshold; the dashed horizontal line indicates the threshold for Z-score *P*-values surviving Bonferroni correction for 7 tested traits. SCZ = schizophrenia; ADHD = attention deficit hyperactivity disorder; BPD = bipolar disorder; MDD = major depressive disorder; ASD = autism spectrum disorder; BLD TG = blood triglyceride levels; HGHT = height.

A limitation of GWAS approaches to complex disorders is that identification of the causal genetic variants underlying associations is often complicated by linkage disequilibrium (resulting in multiple variants at a locus displaying similar levels of association) and incomplete functional characterization of non-coding regions of the genome. To refine potentially functional genetic variants driving genome-wide significant associations with neuropsychiatric disorders in GWAS, we identified SNPs that were located within a bulk fetal frontal cortex OCR, associated with any of the 5 tested neuropsychiatric conditions at genome-wide significance (*P* < 5 × 10^−8^) and in strong linkage disequilibrium (r2 > 0.8) with the GWAS index SNP at the locus. A total of 68 such OCR SNPs were identified for schizophrenia and 15 for bipolar disorder, of which 12 (on chromosome 6) were shared between the two conditions. We further characterize these OCR SNPs in terms of whether they are also in detected fetal brain H3K4me1 and / or H3K4me3 sites (RoadMap Epigenomics Mapping Consortium, 2015), can be confidently attributed to NeuN+ and / or NeuN− nuclei and if they have been found to be a high confidence eQTL (*P* < 5 × 10^−5^) for any transcripts in human fetal brain (O’Brien et al, 2018) (Supplementary Table S5). We found that OCR SNPs on chromosome 6 were not only in strong LD with schizophrenia and bipolar disorder GWAS index SNPs, but also with the most significant eQTL for three transcripts of the *BTN2A1*, *ZSCAN12P1* and *H4C13* genes (r2 with the top eQTL of between 0.88 and 1) in fetal brain. We also observed schizophrenia-associated OCR SNPs on chromosome 8 associated with fetal brain eQTL for *DDHD2* and *FGFR1* transcripts (r2 with top eQTL of 0.85 and 0.95, respectively), and on chromosome 13 associated with fetal brain eQTL for the long noncoding RNAs *LINC01068* and *LINC01038* (r2 with top eQTL of 0.88 and 1, respectively; Figure 7). We did not identify any OCR SNPs in strong LD with the GWAS index SNPs at genome-wide significant loci for ADHD, ASD or major depressive disorder, although we note that the number of such loci in these GWAS is substantially lower than genome-wide significant loci in the utilized bipolar disorder and schizophrenia GWAS.

**Figure 7.**
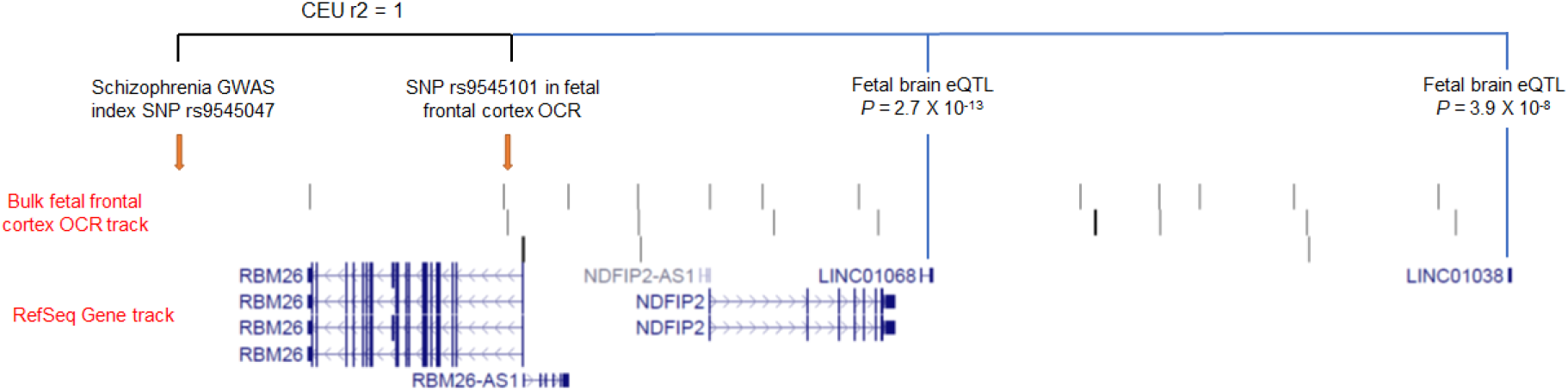
Use of open chromatin region maps to identify potentially functional genetic variants at GWAS loci. In this example, a single nucleotide polymorphism (SNP) within a fetal frontal cortex open chromatin region is in perfect linkage disequilibrium (r2 = 1) with the schizophrenia GWAS index SNP at the locus (rs9545047) and is also a significant eQTL for *LINC01068* and *LINC01038* in the human fetal brain (O’Brien et al, 2018). Image generated using the UCSC Genome Browser (https://genome.ucsc.edu/index.html) and an uploaded BED file for identified bulk fetal frontal cortex open chromatin regions (available through https://doi.org/10.6084/m9.figshare.12302387.v1) as a custom track.

## Discussion

In the absence of clear pathology, but a significant genetic component, insights into the underlying biology of neuropsychiatric disorders are likely to be provided by functional maps of the genome in a variety of human neural cell populations. In the present study, we have mapped regions of open chromatin in bulk tissue, NeuN+ and NeuN− nuclei from the prenatal human frontal cortex. By integrating these maps with other epigenomic data from the human fetal brain (Roadmap Epigenomics Consortium, 2015) and summary statistics from recent large-scale GWAS (Ripke et al, 2020; Wray et al, 2018; Demontis et al, 2019; Grove et al, 2019; Mullins et al, 2021), we provide evidence for an early neurodevelopmental component to a range of neuropsychiatric conditions, and highlight an important role for regulatory regions active within both NeuN+ and NeuN− cells of the prenatal brain in susceptibility to these disorders.

Our findings add to growing evidence that a proportion of the common genetic variants conferring risk to neuropsychiatric disorders are active during prenatal brain development (e.g. Bray & Hill, 2012; Hannon et al, 2016; O’Brien et al, 2018; de la Torre et al, 2018; Li et al, 2018; Schork et al, 2019; Cross-Disorder Group of the Psychiatric Genomics Consortium, 2019; Clifton et al, 2019; Forsyth et al, 2020; Hall et al, 2020). A previous study mapped OCRs in the germinal zone (comprising the ventricular zone, subventricular zone and intermediate zone) and cortical plate (encompassing the subplate, cortical plate and marginal zone) of the human fetal cerebral cortex at 15-17 PCW, reporting significant enrichment of SNP heritability for ADHD, depressive symptoms, neuroticism and schizophrenia within sites that were found to be preferentially accessible in the germinal zone (de la Torre-Ubieta et al, 2018). In the present study, we confirm enrichment of SNP heritability for ADHD, major depressive disorder and schizophrenia within OCRs of the human (frontal) cortex during the second trimester of gestation, and extend this observation to also include bipolar disorder and ASD when either H3K4me1 or H3K4me3 sites were additionally considered.

Although bipolar disorder is not generally considered to be neurodevelopmental in origin, the present data, showing SNP heritability of bipolar disorder to be enriched within fetal brain OCRs at a similar level to that of schizophrenia and ADHD, are consistent with our previous finding of an enrichment of fetal brain eQTL within common genetic risk variants for the condition (O’Brien et al, 2018) and with recent evidence for altered expression of neurodevelopmental genes in iPSC-derived cerebral organoids generated from patients with bipolar disorder (Kathuria et al, 2020). Emerging evidence suggests that some of the common genetic influences on risk for major depressive disorder also operate *in utero* (Wray et al, 2018; Hall et al, 2020), although we note that, in the present study, the enrichment of SNP heritability for the condition in fetal brain OCRs was consistently lower than that for bipolar disorder and schizophrenia. While rare genetic risk variants for ASD are known to disrupt genes functioning in the prenatal brain (Willsey et al, 2013; Parikshak et al, 2013; Satterstrom et al, 2020), only recently have ASD GWAS yielded sufficient signal for biological insights into the condition (Grove et al, 2019, Pain et al, 2019; Forsyth et al, 2020; Hall et al, 2020).

Our study provides the first maps of open chromatin in NeuN+ and NeuN− cell populations of the human fetal brain. In contrast to the adult brain, where genetic risk for schizophrenia appears to be largely mediated by NeuN+ cells (Fullard et al, 2018), we find that, within the fetal brain, genetic risk for this and other neuropsychiatric conditions is at least as strongly enriched within high confidence OCRs observed in NeuN− nuclei. However, whereas NeuN− cells in the adult brain largely consist of mature glia (astrocytes and oligodendrocytes), those of the second trimester fetal brain encompass a variety of developing cells, including intermediate progenitors, radial glia and oligodendrocyte precursors. Our data are therefore consistent with the findings of de la Torre-Ubieta and colleagues (2018), who report enrichment of SNP heritability for neuropsychiatric disorders in OCRs of the neural progenitor cell-containing germinal zone, and of Schork and colleagues (2019), who report enriched expression of fine-mapped candidates genes for a broad neuropsychiatric phenotype in fetal radial glia. However, it is possible that the NeuN− nuclei assayed in this study also include those from neurons at early stages of development. Mapping of OCRs in human prenatal brain using single cell / nuclei sequencing technologies will be necessary to elucidate the specific fetal cell populations relevant to neuropsychiatric disorders.

We used our high confidence fetal frontal cortex OCRs to identify potentially functional non-coding SNPs tagged by genome-wide significant risk SNPs for schizophrenia and bipolar disorder, which may serve as a guide for future functional studies (e.g. genome editing). Although we have focused on neuropsychiatric phenotypes, our maps of open chromatin in the fetal frontal cortex are likely to be useful in exploring early neurodevelopmental antecedents to a variety of brain-related conditions. As we move into the era of whole genome sequencing, maps of functional elements within the non-coding genome will continue to be important. We therefore provide the genomic coordinates of our high confidence fetal brain OCRs through a publicly-accessible online repository (https://doi.org/10.6084/m9.figshare.12302387.v1) for use by the research community.

## Acknowledgements

This work was supported by a Medical Research Council (MRC) GW4 BioMed Doctoral Training Partnership studentship to MRK, an MRC project grant to NJB (grant number MR/L010674/2) and an MRC Centre grant (grant number MR/L010305/1). The human fetal material was provided by the Joint MRC/Wellcome Trust (grant #099175/Z/12/Z) Human Developmental Biology Resource (www.hdbr.org). We thank Dr Joe Burrage, Dr Catherine Naseriyan and Dr Ann Kift-Morgan for assistance with fluorescence-activated nuclei sorting, and Dr Joanne Morgan for assistance with sequencing. We thank the schizophrenia and bipolar disorder working groups of the Psychiatric Genomics Consortium for use of summary GWAS data prior to publication. We thank the research participants and employees of 23andMe for their contribution to the major depressive disorder GWAS data used in this study. Comparison adult frontal cortex ATAC-Seq data were generated as part of the CommonMind Consortium, supported by funding from Takeda Pharmaceuticals Company Limited, F. Hoffmann-La Roche Ltd and NIH grants R01MH085542, R01MH093725, P50MH066392, P50MH080405, R01MH097276, RO1-MH-075916, P50M096891, P50MH084053S1, R37MH057881, AG02219, AG05138, MH06692, R01MH110921, R01MH109677, R01MH109897, U01MH103392, and contract HHSN271201300031C through IRP NIMH. CMC Leadership: Panos Roussos, Joseph Buxbaum, Andrew Chess, Schahram Akbarian, Vahram Haroutunian (Icahn School of Medicine at Mount Sinai), Bernie Devlin, David Lewis (University of Pittsburgh), Raquel Gur, Chang-Gyu Hahn (University of Pennsylvania), Enrico Domenici (University of Trento), Mette A. Peters, Solveig Sieberts (Sage Bionetworks), Thomas Lehner, Stefano Marenco, Barbara K. Lipska (NIMH).

## Data Availability Statement

The genomic coordinates of the fetal frontal cortex open chromatin regions defined and used in this study are provided as BED files in a publicly-accessible online repository: https://doi.org/10.6084/m9.figshare.12302387.v1

