## Supplementary Figures for "Sites of active gene regulation in the prenatal frontal cortex and their role in neuropsychiatric disorders"

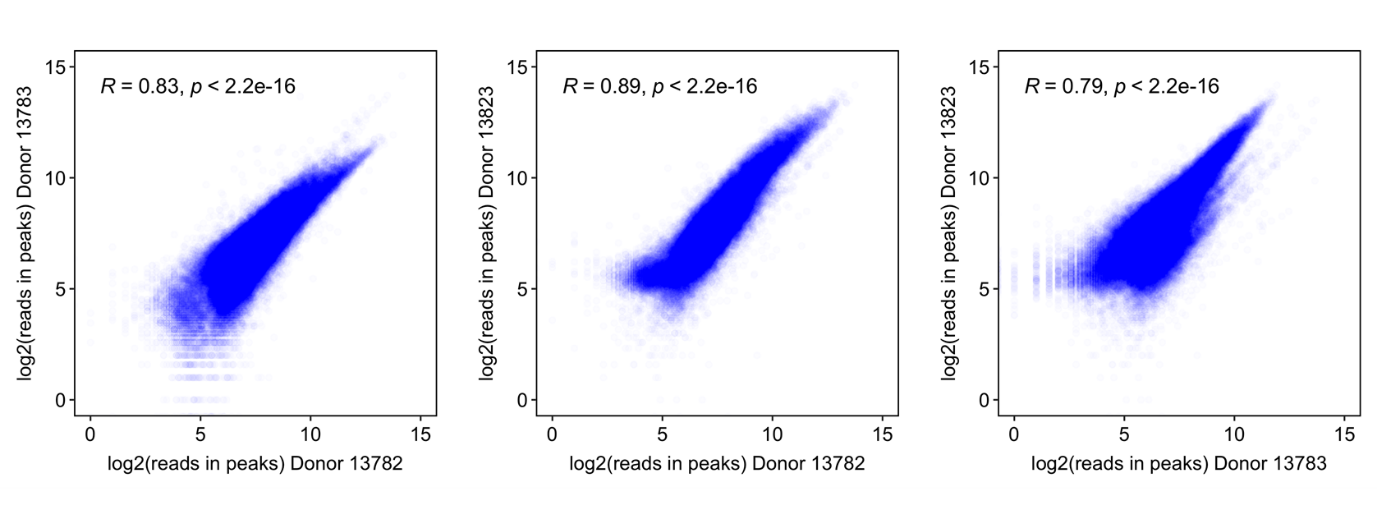

**Supplementary Figure S1.** Correlations between the 3 donors in sequencing reads within high confidence bulk fetal frontal cortex open chromatin regions.

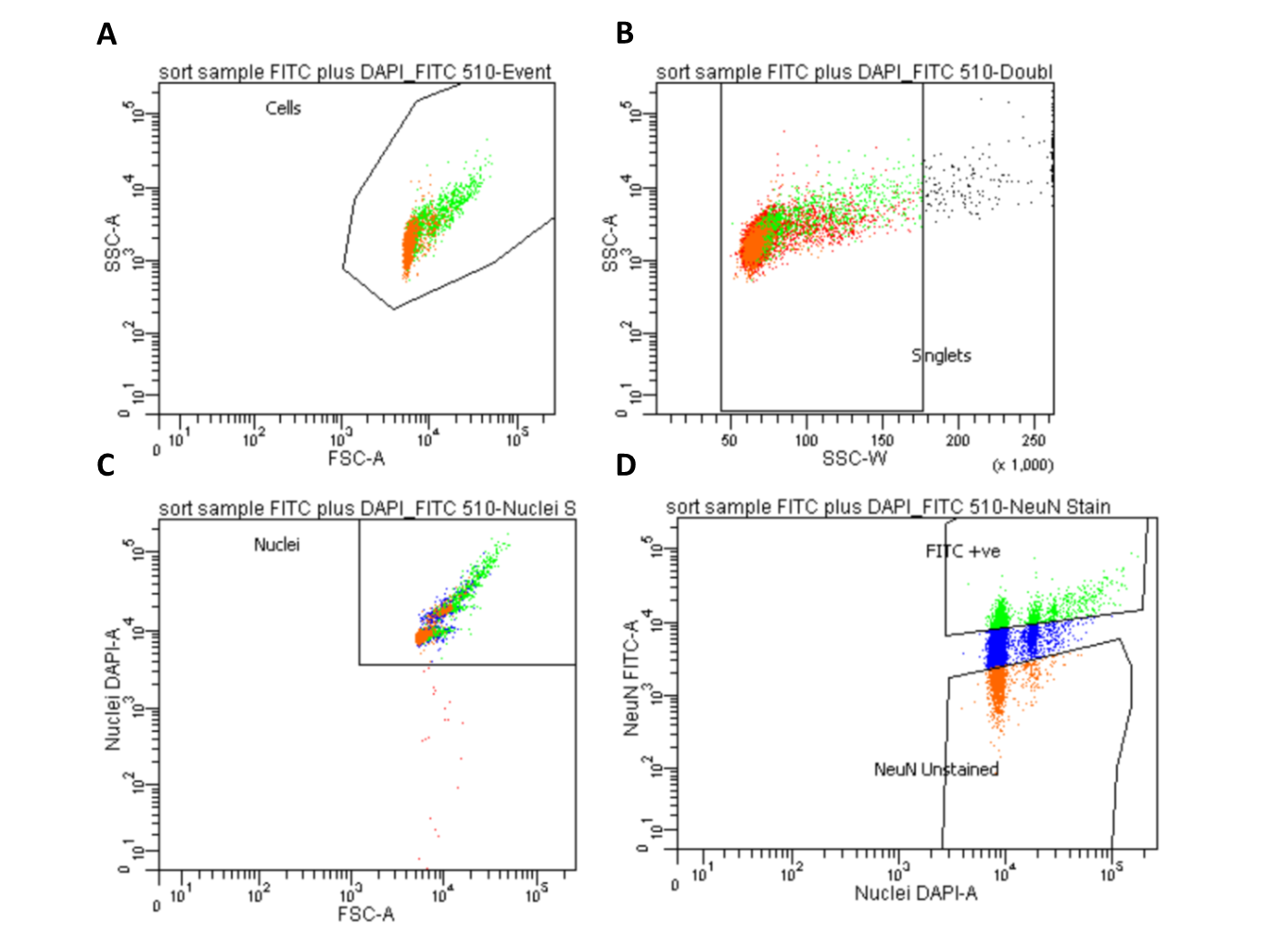

**Supplementary Figure S2. Isolation of NeuN+ and NeuN- nuclei using fluorescence-activated nuclei sorting (FANS).** First, debris was excluded using forward and side scatter pulse area parameters (FSC-A and SSC-A) (A). This was followed by exclusion of nuclei aggregates using pulse width (FSC-W and SSC-A) (B) and isolation of DAPI-positive nuclei (C), before gating populations based on NeuN fluorescence (D).
